# Trait anxiety is associated with idiosyncratic neural event boundaries during naturalistic movie-watching

**DOI:** 10.64898/2026.01.03.697513

**Authors:** Alicia T.H. Liu, Jiajie Chen, Jadyn Park, Haena Kim, Yuan Chang Leong

## Abstract

Anxiety is associated with altered patterns of attention to images and objects, but how it influences the perception of continuous experiences remains underexplored. Here, we examined whether trait anxiety is associated with differences in how the brain segments continuous experience into events. Using a large-scale open-access dataset and data-driven methods to detect neural event boundaries, we compared neural event segmentation between healthy adults with high (N=60) and low trait-level anxiety (N=60) as they viewed a short suspenseful film. While the cortical temporal hierarchy was preserved across groups, high-anxiety individuals had more idiosyncratic event boundaries, exhibiting reduced alignment to both low-anxiety individuals and one another. This divergence was most pronounced in the dorsal attention and frontoparietal control networks, suggesting that trait anxiety is associated with idiosyncratic temporal organization in systems involved in attentional control. Furthermore, boundary variability in high-anxiety participants was greater during moments rated as more anxiety-provoking by a large language model, controlling for arousal, valence, and low-level visual features. Together, these findings suggest that anxious individuals organize continuous experience in a more individualized manner. More broadly, they offer a neurocomputational framework for investigating how trait-level variability interacts with narrative content to shape naturalistic event perception.

## 1. Introduction

Anxiety-related disorders are among the most prevalent class of mental health conditions, affecting over 300 million people as the sixth leading cause of disability worldwide (Yang et al., 2021). Individuals with high trait anxiety experience persistent fear and worry related to potential threats (Heeren et al., 2018). Research links this to attentional biases towards negative stimuli, hyperactivity in the amygdala, insula, and anterior cingulate, and differences in white matter connectivity between the amygdala and the prefrontal cortex (Bar-Haim et al., 2007; Bishop, 2009; Kim & Kim, 2022; Sylvester et al., 2012). However, it remains unclear how anxiety shapes the neural organization of experiences over time: everyday experiences are not composed of isolated stimuli, but of continuous streams of information. Does anxiety also alter the way we segment this flow into events, impacting how daily experiences are perceived?

The brain is thought to organize continuous experience into chunks of meaningful information to efficiently encode them into memory (Zacks & Swallow, 2007; Zwaan et al., 1995). This segmentation plays a fundamental role in how we perceive, interpret, and remember ongoing experience (Zacks, 2020). Effective segmentation supports comprehension and recall, while poor segmentation leads to fragmented or distorted memory (Sargent et al., 2013; Swallow et al., 2009; Zacks et al., 2006). Event segmentation theory proposes that segmentation depends on event models, which are multimodal representations combining external input with schematic knowledge to predict what will happen next (Kurby & Zacks, 2008; Zacks et al., 2007). For example, when at a restaurant, we form an event model for ‘dining out’. This event model enables predictions of upcoming actions such as eating the meal and asking for the check.

Recent neuroimaging work shows that the brain segments continuous experiences by transitioning between stable neural states, with neural event boundaries aligning with behaviorally identified event boundaries (Baldassano et al., 2017; Geerligs et al., 2022). These neural boundaries are organized hierarchically, with sensory regions representing short, rapidly changing events, while higher-order regions integrate information over longer timescales. Neural event segmentation has been applied where behavioral segmentation is not feasible, such as in infants, revealing fewer, longer events and less hierarchical differentiation compared to adults (Yates et al., 2022). Importantly, neural event boundaries are not entirely determined by stimulus content, but also shaped by individual traits such as goals, attention, and schematic knowledge (Soares et al., 2024). This individual variability in neural event segmentation has been linked to behavior, with participants whose neural event boundaries are more aligned during narrative processing exhibiting similar interpretations and memories of the narrative (Sava-Segal et al., 2023).

How individual traits such as anxiety affect neural event segmentation remains unclear. Cognitive models of anxiety propose systematic biases in attention that shape expectations about the environment and influence goal selection (Eysenck et al., 2007; Mathews & MacLeod, 2005). These biases could alter the schematic knowledge that individuals bring to bear when updating event models. For example, a non-anxious person might interpret a waiter’s glance as neutral, while an anxious person might infer irritation, altering subsequent expectations. Such shifts in interpretation and expectations could change where event boundaries are inferred. As a result, the same experience may be segmented into different units by anxious individuals (Sherrill et al., 2019).

Here, we investigate how trait anxiety influences the neural segmentation of continuous experience using an open-access fMRI dataset of participants watching a suspenseful film (Taylor et al., 2017). We applied data-driven methods to detect neural event boundaries across cortical regions, and asked whether anxious and non-anxious individuals differed in the alignment of neural event boundaries. We utilized a continuous metric of distance to quantify boundary alignment, giving us a fine-grained measure of individual differences in event segmentation. We used a large language model to generate affective ratings from moment-to-moment annotations of the movie and validated these ratings against human ratings in an independent sample. This allowed us to ask whether anxiety-related differences in neural event segmentation were linked to the affective content of the unfolding narrative. Together, this work characterizes how trait anxiety relates to the temporal organization of continuous experience in the brain and offers a framework for investigating individual differences in naturalistic event cognition.

## 2. Materials & Methods

### 2.1 Ethics Statement

Data analysis was approved by the Institutional Review Board of the University of Chicago (IRB22-0002). All data were anonymized by the original authors. Collection of data was approved by the Cambridgeshire 2 Central Ethics Committee (reference: 10/H0308/50). Written informed consent was obtained from all participants.

### 2.2 Participants

Data used in the preparation of this work was obtained from the Cambridge Centre for Ageing and Neuroscience data repository (available at https://opendata.mrc-cbu.cam.ac.uk/projects/camcan/) (Shafto et al., 2014; Taylor et al., 2017). The full sample consisted of 623 participants (316 female), from which 9 participants’ data were excluded due to imaging quality issues. All participants were native English speakers, physically healthy, had no current drug abuse, no history of serious head injury, or current serious psychiatric conditions such as bipolar disorder or schizophrenia. For full exclusion criteria, see Table 1 in Shafto et al. (2014).

From this sample, we sought to identify two equally-sized control and anxiety groups. We assessed trait anxiety based on the anxiety subscale scores of the Hospital Anxiety and Depression Scale (HADS), which consists of 7 items (Zigmond & Snaith, 1983). Each item is scored on a 4-point scale (0-3), with the subscale total ranging from 0 to 21; HADS-A scores of 0-7 indicate normal, 8-10 mild, 11-14 moderate, and 15-21 severe anxiety. We excluded participants above the age of 65 as prior studies found age-related changes in event boundary-related neural activity (Lugtmeijer et al., 2025; Reagh et al., 2020). Among the remaining participants, anxiety scores ranged from 0 to 19 (M = 6.54, SD = 4.19).

Out of the 363 remaining participants, 60 individuals had mild to severe anxiety (HADS-A of >7), and these participants were designated as the anxious group. The 60 participants with the lowest anxiety scores were then selected to form the non-anxious group. The anxious group had a mean score of 10.2 (SD=2.35), while the non-anxious group had a mean score of 2.75 (SD=1.25). The two groups did not differ in age (anxiety M = 34.7, SD = 7.90; healthy M = 35.6, SD = 7.97) or sex composition (Mann-Whitney U test, p = 0.423; Chi-squared test, p = 0.715). The two groups were also matched on cognitive battery scores including word memory, verbal fluency, immediate logical memory, and delayed logical memory (Mann-Whitney U test p > 0.05). During quality control checks, one participant in the non-anxious group was excluded due to a verbal fluency score more than 3 standard deviations below both group means (score = 6, anxiety M = 12.45, SD = 1.76, healthy M = 12.07, SD = 1.98).

### 2.3 Stimulus

The audiovisual stimulus is a condensed version of drama ‘Bang! You’re Dead’ by Alfred Hitchcock. The original version is 25 min; the version presented to participants was shortened to 8 min 13 s and preserves the narrative arc and flow of the film (Shafto et al., 2014). This abbreviated version of ‘Bang! You’re Dead’ has been shown to be effective in evoking reliable time locked brain activity across participants (Hasson et al., 2004) and has also been used to study neural event segmentation in prior studies (Lugtmeijer et al., 2025; Reagh et al., 2020).

### 2.4 fMRI data acquisition & pre-processing

All MRI data was collected by Cam-CAN at the UK Medical Research Council Cognition and Brain Sciences Unit, based in Cambridge, England. A multi-echo T2*-weighted gradient-echo EPI sequence with a 3T Siemens TIM Trio scanner and a 32-channel head coil was used. Each scan contained 193 volumes, with 32 axial slices per volume, each 3.7 mm thick and separated by a 0.74 mm gap. The repetition time (TR) was 2470 ms, and five echo times (TE) were used: 9.4 ms, 21.2 ms, 33 ms, 45 ms, and 57 ms. The flip angle was 78°, with a field of view (FOV) of 192 mm × 192 mm and voxel dimensions of 3 mm × 3 mm × 4.44 mm. The total acquisition time was 8 minutes and 13 seconds. The first TR corresponded to the first frame of the movie stimulus.

Data underwent pre-processing in the standard Cam-CAN pipeline (Taylor et al., 2017). The functional data was first combined across five echoes. In AFNI (version AFNI_17.1.01; https://afni.nimh.nih.gov; Cox, 1996), the resulting time series were unwarped using field-map images to correct for magnetic field inhomogeneities, followed by slice-time correction, and realignment to correct for head motion. Then, using SPM 12 software (http://www.fil.ion.ucl.ac.uk/spm; Ashburner, 2007), data was co-registered to T1-weighted structural scans, DARTEL intersubject alignment was performed, and normalized to MNI space using transformation parameters derived from the structural preprocessing pipeline.

In addition to the Cam-CAN project pre-processing, we applied a high-pass filter (cutoff = 0.008 Hz) to remove slow signal drifts. We then parcellated the brain into 200 cortical regions of interest (ROI) using the Schaefer-200 atlas with Yeo 7-network solution (Schaefer et al., 2018), removed voxels with zero variance across time and z-scored and averaged data within each region.

### 2.5 Characterizing temporal cortical hierarchy in anxious and non-anxious groups

We first fit a Shared Response Model (SRM) with 40 features separately to the anxious and non-anxious group ROI timecourses using the BrainIAK toolkit (Kumar et al., 2021). The SRM projects individual participant data into a shared feature space, emphasizing variance that is consistent across participants while reducing noise from idiosyncratic activity patterns. This step improves our ability to detect reliable group-level effects by aligning functional data across individuals (P.-H. (Cameron) Chen et al., 2015).

We then applied Greedy State Boundary Search (GSBS) to identify the number of events separately for anxiety and non-anxious groups (Geerligs et al., 2021). We chose to use GSBS to estimate group-level event number as it has been shown to operate reliably at a group size of greater than 20 participants and outperforms Hidden Markov Model (HMM) methods in terms of computational speed. GSBS uses a greedy search algorithm that iteratively places boundaries which maximize within-event voxel activity pattern similarity. Similarly, it estimates the optimal number of events by testing a potential range and selecting that which maximizes within-event pairwise voxel activity correlation and minimizes pairwise voxel activity correlation between neighboring events.

To assess whether the cortical hierarchy is preserved between the two groups, we computed the Spearman correlation between the number of events per ROI of the two groups. We also examined regions previously associated with shorter-versus longer-timescale events: primary and extrastriate visual areas, intraparietal sulcus, angular gyrus, dorsal medial prefrontal cortex, and ventromedial prefrontal cortex, expecting the most events in primary visual cortex and the fewest in prefrontal regions (Baldassano et al., 2017; Geerligs et al., 2022). We defined these regions by averaging event counts across Schaefer-200 ROIs that overlapped with activation maps from NeuroSynth (https://neurosynth.org/), an open platform that aggregates data from fMRI studies to generate probabilistic brain activation maps for specific terms. Specifically, for each target region (e.g., V1), we downloaded the corresponding NeuroSynth activation map, visualized it alongside the Schaefer-200 parcellation in FSLeyes (https://pypi.org/project/fsleyes/), identified Schaefer parcels falling within the NeuroSynth-defined region, and averaged their event counts.

We formally evaluated group differences in event segmentation rate (i.e., the number of events per region of interest) with non-parametric permutation testing. Group labels were shuffled and event numbers were recomputed with GSBS 5,000 times to generate a null distribution. We used an alpha level of 0.05 as the threshold for statistical significance and applied false discovery rate correction with the Benjamini-Hochberg procedure (Benjamini & Hochberg, 1995).

### 2.6 Detecting event boundaries

We build on recent work demonstrating that Hidden Markov Models (HMMs) can reliably detect event boundaries at the individual level (Sava-Segal et al., 2023). Similar to GSBS, HMMs segment the time course into a sequence of discrete neural states, each characterized by a stable multivariate pattern of voxel activity, with state transitions defining event boundaries (Baldassano et al., 2017). To obtain a common event number for each ROI across both groups, we re-fit GSBS to data across all participants. This ensures that boundary alignment comparisons were not biased by differences in event counts between groups. We chose HMMs for individual-level event boundary estimation as existing studies find that they reliably estimate participant-level event boundaries (Antony et al., 2021; Sava-Segal et al., 2023; Lu et al., 2025; Zhou et al., 2026), while GSBS is validated on and more commonly used with group-averaged data (Geerligs et al., 2021; Oetringer et al., 2025). In contrast, we used GSBS rather than HMMs for event-number estimation because GSBS has been shown to recover the number of events more accurately than HMM-based approaches while also requiring substantially less computational time (Geerligs et al., 2021).

With the per-ROI event number estimated by GSBS, we fit a HMM to each individual’s time series. As we were interested in preserving individual idiosyncrasies in the neural time series and the model was fit to each individual separately, we did not transform the time series with the SRM when fitting the HMM. The HMM fitting produced neural event boundary locations for every ROI in each participant.

### 2.7 Computing neural boundary alignment with Earth Mover’s Distance

To examine anxiety-related differences in event boundary patterns, we computed a pairwise neural alignment metric for each ROI. For each participant pair, we identified the event boundary timepoints estimated by the HMM and measured the dissimilarity between their boundary patterns using Earth Mover’s Distance (EMD), as implemented in SciPy (https://docs.scipy.org/doc/). EMD is a measure of similarity between two probability distributions, conceptualized as the minimum cost of transforming one distribution into the other (Rubner et al., 2000).

Intuitively, EMD quantifies how much “work” it takes to transform one set of event boundaries into another, treating each as a distribution over time. The more similar two participants’ boundary patterns are, the less work is needed to align them. In our study, the number of events in each ROI was fixed across participants, so EMD reduces to the average displacement (timing difference) between order-matched boundary pairs. We adopt the EMD framework because it provides a principled and unified metric that generalizes to cases with differing numbers of boundaries, such as comparisons across brain regions or participant groups with different event counts. Compared to methods that count coinciding boundaries within a fixed time window (Baldassano et al., 2017; Sava-Segal et al., 2023; Soares et al., 2024), EMD provides a more graded measure of temporal alignment between event boundary sets as it considers not just whether two boundaries match within a fixed window but also how far they are apart. This yielded one EMD score per participant pair per ROI, with higher values indicating greater dissimilarity in event boundary timing. As a robustness check, we repeated our analysis using the boundary-match counting method, considering boundaries occurring within 1 TR (2.47 s) as a match.

For each ROI, we fit Bayesian multilevel models to model EMD score as a function of dyad composition (anxious-anxious, non-anxious-non-anxious, anxious-non-anxious) (G. Chen et al., 2020). To account for the dependence structure arising from each participant contributing to multiple dyads, random effect intercepts for each member of the dyad were included in each model. We performed three planned contrasts: (1) anxious-anxious > non-anxious-non-anxious, (2) anxious-anxious > anxious-non-anxious, and (3) anxious-non-anxious > non-anxious-non-anxious to test the hypothesis that boundary alignment was more dissimilar (corresponding to higher EMD) in dyads that included more anxious individuals. For each contrast, we computed the proportion of the posterior distribution greater than 0 as the estimate of the probability of a credible effect.

We note that anxiety scores were more variable in the anxious group (SD = 2.35) than in the non-anxious group (SD = 1.25). Thus, an alternative explanation for higher EMD in anxious dyads is that it reflects larger score differences within those dyads, rather than idiosyncrasy associated with trait anxiety. To address this possibility, we ran an intersubject representational analysis (Finn et al., 2020) using a continuous measure of dyadic anxiety. We computed the mean anxiety score of each dyad, and used Bayesian multilevel models to test its relationship with dyadic EMD score. Dyads with the highest anxiety scores would have a high mean but a small score difference. Thus, if higher mean anxiety scores tracked with higher EMD scores, it would indicate that event boundary divergence scaled with anxiety levels rather than the difference in anxiety scores. Models again included random effect intercepts for each member of the dyad.

To account for multiple comparisons across ROIs, we thresholded the brain maps at an expected posterior error probability (PEP) of 0.01, corresponding to a less than 0.01 probability of making an incorrect inference (Käll et al., 2008; Storey, 2003). From a frequentist perspective, this approach is analogous to controlling for a false discovery rate at q < 0.01. Models were fit using the brms package in R (Bürkner, 2017) with MCMC sampling (4 chains, 1000 warm up, 2000 iterations each) using weakly informative priors (Student’s t(3, 0, 1) for random effect standard deviations and half-Cauchy(0, 1) for residual variance). Model convergence metrics indicated adequate sampling across ROIs (average maximum R-hat = 1.02; average minimum ESS ratio = 0.08), yielding a minimum ESS of approximately 320 samples, above suggested minimum values for reliable estimates (G. Chen et al., 2020).

### 2.8 Extracting affective and visual features from movie

Building on previous work showing that large language models can accurately detect emotional states and affective experience from text annotations (Park et al., 2025; Rathje et al., 2024) and detect narrative structure with human-level performance (Michelmann et al., 2025; Panela et al., 2025), we leveraged a large language model, GPT-4o, to generate scene-level affective ratings of the movie stimulus. We divided the movie into 4-TR (9.88 s) segments, producing 48 segments from the 8-minute film. Following procedures described in previous work (J. Chen et al., 2017; Lee & Chen, 2022), we manually annotated these segments with a brief text description of the events occurring onscreen and included direct quotes of any spoken dialogue. When dialogue spanned multiple segments, it was split across segment annotations so that each portion of the quote appeared only in the segment in which it occurred.

For each segment, we provided the current and all prior annotations to GPT-4o and instructed the model to rate the current segment on a scale of 1-7 along one of three dimensions: anxiety, arousal, and valence. Anxiety was the primary dimension of interest, while arousal and valence were collected as covariates to test the specificity of anxiety-related effects. Ratings for each dimension were obtained separately to prevent ratings on one dimension from influencing ratings on the others. We set the model temperature to 0 for deterministic rating. We used the system prompt “You are an expert movie rater, rating scenes from a movie as it unfolds” and provided GPT-4o with the definition of the dimension being rated:

● *“Anxiety refers to feeling tense, nervous, or apprehensive about what may happen, with 1 being calm and at ease and 7 being worried and on edge.”*
● *“Arousal refers to feeling very mentally or physiologically alert, activated and/or energized, with 1 being drowsy and sluggish and 7 being highly alert and energized.”*
● *“Valence refers to how positive or negative the emotional experience feels, with 1 being very unpleasant and negative and 7 being very pleasant and positive.”*

The definition of anxiety was based on the definition provided to human participants in our validation dataset (Morgenroth et al., 2025; see below) and aligns with established conceptualizations of anxiety as anticipatory apprehension about uncertain future events (Grupe & Nitschke, 2013). The definitions of valence and arousal were drawn from our earlier work (Ke et al., 2025; Pan et al., 2025; Park et al., 2025). In addition to the GPT-4o ratings, we used OpenCV (https://opencv.org/) to extract low-level visual features (brightness, motion, and optical flow) every 5 frames in the movie and averaged these features within 4-TR windows to include them as covariates in our model.

We validated the GPT-4o anxiety ratings using the Emo-FilM dataset (Morgenroth et al., 2025) which contains moment-by-moment affective ratings of 16 short films on multiple affective dimensions, including anxiety, from 44 human participants. We selected three films from the dataset that spanned different genres and were similar in length to *Bang! You’re Dead* (see *Supplementary Materials*). The films were manually annotated using the same procedure, and the resulting segment-by-segment annotations were submitted to GPT-4o using the same prompt, rating scales, and model settings.

LLM-generated arousal and valence ratings have previously been found to correlate with human ratings in naturalistic datasets (arousal: Pan et al., 2025; Park et al., 2025; valence: Yang et al., 2024). Here, we similarly validated our GPT-4o arousal and valence ratings against human ratings. As arousal and valence ratings were not available for the Emo-FilM dataset, we correlated GPT-4o arousal and valence ratings with related dimensions of alertness and happiness, respectively. As a more direct validation, we compared the LLM-generated arousal and valence ratings against human ratings of arousal and valence for a 48-min segment from an episode of BBC’s *Sherlock* obtained from 30 participants in a previous study (Ke et al., 2025).

For each affective dimension, GPT-4o ratings were correlated with average ratings across human participants using Pearson correlation. Statistical significance was assessed with a phase randomization permutation test, a nonparametric method that generates surrogate timeseries preserving the original power spectrum and autocorrelation structure while disrupting temporal alignment (Lancaster et al., 2018; Theiler et al., 1992). We generated 5,000 phase-randomized surrogates of the GPT-4o ratings and computed the correlation between each surrogate and the average human participant ratings to construct a null distribution. Two-tailed *p*-values were computed as the proportion of null correlations equal to or exceeding the observed *r* in absolute value.

### 2.9 Relating movie features to idiosyncratic event boundaries

We tested whether event boundaries were more idiosyncratic in anxious individuals than non-anxious individuals during anxiety-inducing moments in the movie. We restricted our analysis to ROIs that showed significantly higher EMD in anxious dyads than non-anxious dyads identified in the earlier analyses. For each ROI and each participant, the HMM yielded K event boundaries ordered chronologically, where K is the GSBS-derived number of events. For each boundary position i (i = 1, 2, … K), we computed the variance in boundary timing across participants separately within the anxious and non-anxious groups, yielding two variance values per boundary. This captures how temporally clustered or dispersed each boundary position was across participants within the anxious and non-anxious groups.

We then computed a variance ratio score for each boundary:

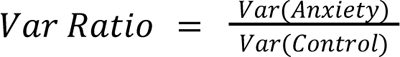

refer to variance in boundary locations for anxious individuals and non-anxious individuals, respectively. This ratio reflects how much more temporally variable anxious individuals were compared to non-anxious individuals at this boundary. We used the ratio rather than the raw difference to ensure comparability across ROIs with different baseline number of events.

To temporally align neural event boundaries with stimulus content, we shifted the BOLD timecourses by 2-TRs (4.94s) to account for hemodynamic lag. For each boundary, we computed the average boundary time across participants, which provided a single best estimate of when that boundary occurred in the movie. To assess the overall relationship between on-screen stimulus features and boundary variability, we fit a Bayesian multilevel model to predict variance ratio across all ROIs from the average anxiety rating of the 4-TR segments that fell within the HMM-derived event immediately preceding each boundary. Segments were included only if they fell fully within the event. For instance, if a boundary occurred at TR 40 and the preceding event spanned TRs 25-39, we averaged anxiety ratings of the segments that are entirely within TRs 25-39.

ROIs were modeled as random effects, allowing us to model the dependence within each ROI and take advantage of partial pooling across ROIs (G. Chen et al., 2019). As event boundaries were specific to each ROI, the set of segments associated with a given boundary varied across ROIs. Arousal ratings, valence ratings, and low-level features were similarly averaged in the preceding event and included in the model as covariates. We used weakly informative priors Normal(0,1) for the intercept and fixed effects, and exponential (1) for random effect standard deviations and residual variance.

### 2.10 Decoding neural signatures of sustained attention during movie-watching

To assess group differences in sustained attention to the stimulus, we applied the sustained attention connectome-based predictive model (saCPM; Rosenberg et al., 2016) to each participant’s functional data. The saCPM is a validated functional connectivity model that predicts sustained attention during cognitive tasks from whole-brain functional connectivity patterns (Rosenberg et al., 2016, 2020), and has been shown to generalize to predict narrative engagement to movies and audio stories (Corriveau et al., 2026; Song et al., 2021). The saCPM consists of two networks of edges (high-attention and low-attention) whose strengths positively and negatively predict sustained attention, respectively. We applied the high- and low-attention networks defined in Song et al. (2021) to each participant’s data. Following Song et al. (2021), each participant’s data was parcellated using the 114-region Yeo cortical atlas combined with 8 subcortical regions from the Brainnetome atlas. We computed dynamic functional connectivity using a sliding-window approach (window = 20 TR, Gaussian sigma = 3 TR), Fisher z-transforming the resulting Pearson correlations. At each timepoint, a decoded sustained attention score was computed as the mean FC strength across positive-network edges minus that of negative-network edges, with higher scores reflecting a connectivity pattern associated with higher sustained attention. Each participant’s resulting decoded attention time course was then averaged within each 4-TR segment and z-scored within participant. This yielded a segment-by-segment measure of decoded attention.

To compare decoded sustained attention between high- and low-anxiety moments, we divided segments based on a tertile split of GPT-4o anxiety ratings (high-anxiety: top tertile; low-anxiety: bottom tertile). Robustness checks were performed using median and quartile splits. Within each group, decoded attention scores for high- versus low-anxiety segments were compared using paired t-tests. To test for group differences in this contrast, we fit a linear mixed-effects model with decoded sustained attention score as the outcome, and segment type (high vs. low anxiety), group (anxious vs. non-anxious), and their interaction as fixed effects, with participant as a random intercept. The interaction term tested whether the effect of segment anxiety on sustained attention differed between groups. As a robustness check, we repeated the analysis with the anxiety rating of each segment as a continuous predictor, modelling between-subject variation in the relationship between moment-level anxiety and attention as a random slope over participants.

## 3. Results

### 3.1 Temporal cortical hierarchy is consistent across anxious and non-anxious individuals

Participants in the anxious group exhibited a mean HADS-A score of 10.2 (SD=2.35), while those in the non-anxious group had a mean score of 2.75 (SD=1.25) (Figure 1A). Prior work found that neural event segmentation exhibits hierarchical organization characterized by a posterior-to-anterior gradient from shorter events in early sensory regions to longer events in higher-order association areas, with brain regions subserving similar cognitive functions demonstrating similar number of events (Baldassano et al., 2017; Honey et al., 2012; Sava-Segal et al., 2023). We first assessed the extent to which this temporal cortical hierarchy was preserved across anxious and non-anxious individuals.

**Figure 1:**
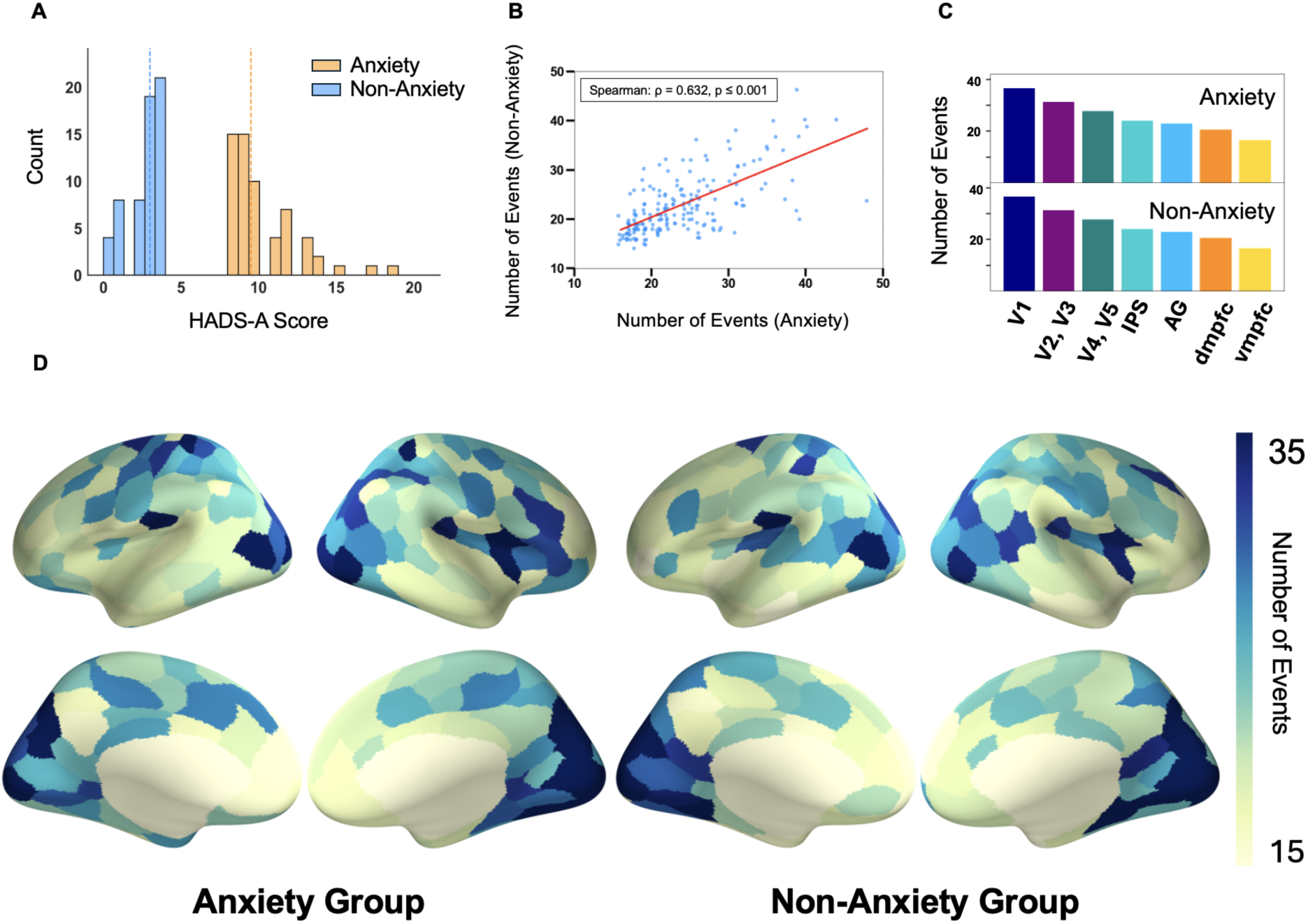
Both anxious and non-anxious groups displayed the expected hierarchy of cortical events. **(A)** HADS-A score distribution of the two groups **(B)** The number of events in each ROI was correlated across the two groups. **(C)** Number of events in ROIs at different stages of the cortical hierarchy, with a clear gradient from sensory to higher order regions across both groups. **(D)** Cortical maps showing the number of neural events identified by GSBS in each ROI. In both groups, the number of events detected decreases along a posterior-to-anterior gradient.

Using GSBS, we estimated the number of events separately for the anxious and non-anxious groups in each ROI. The number of events per region was correlated between the anxious and non-anxious groups (Spearman’s ρ = 0.632, p ≤ 0.001; Figure 1B). When examining selected individual regions, both groups displayed the expected hierarchy, with the most events in the early visual cortex (V1) and the fewest in medial prefrontal cortex (Figure 1C). Finally, when examining the cortical maps, this organization was apparent as a clear posterior-anterior gradient in event segmentation rate, with shorter events in sensory regions and longer events in higher-order association areas (Figure 1D). We then directly compared the event segmentation rates in the two groups within each ROI with permutation testing. Specifically, we compared the observed event segmentation rate to a null distribution generated by shuffling group labels and recomputing per ROI event numbers across 5,000 permutations. There were no ROIs where event segmentation rates were significantly different between groups (FDR-corrected q > 0.05). Together, these results demonstrate that event numbers estimated by GSBS recovered the expected temporal cortical hierarchy of event timescales in both groups.

### 3.2 Anxious individuals exhibit idiosyncratic event boundary patterns

We next examined whether individuals with higher anxiety exhibit idiosyncratic patterns of neural event segmentation. To ensure that measures of boundary alignment were not biased by groups having different numbers of events, we re-fit GSBS across all participants, yielding a single event number per ROI. As was the case when GSBS was fit to the two groups separately, fitting GSBS across both groups reproduced the expected temporal hierarchy in segmentation rates (Figure S1). We then fit an HMM to each individual’s time course using this common event number, which allowed for a fair comparison of boundary locations across groups.

We compared dyadic neural boundary alignment between groups to test whether anxious individuals were less aligned than non-anxious individuals (Figure 2A). For each pair of participants, we computed the dyadic boundary alignment as the earth mover’s distance (EMD) between boundary locations. Using Bayesian multilevel models, we found that EMD was credibly higher between pairs of anxious individuals than between pairs of non-anxious individuals in the lateral and dorsomedial prefrontal cortex, premotor cortex and frontal eye fields, cingulo-opercular regions, intraparietal sulcus, supramarginal gyrus, angular gyrus, superior temporal gyrus, inferior temporal gyrus, temporal pole, and visual cortex (Figure 2B; Table S1). No regions exhibited a credible effect in the opposite direction. Comparable results were obtained when using the number of boundary matches as the measure of boundary alignment, with anxious-anxious pairs having fewer boundary matches than non-anxious-non-anxious pairs (Figure S2). Consistent with greater idiosyncrasy in event segmentation in anxious participants, secondary contrasts revealed that EMD was also higher between anxious-anxious pairs than anxious-non-anxious pairs, and between anxious-non-anxious pairs than non-anxious-non-anxious pairs (Figure S3). As was the case with our main contrast, no regions exhibited a credible effect in the opposite direction.

**Figure 2:**
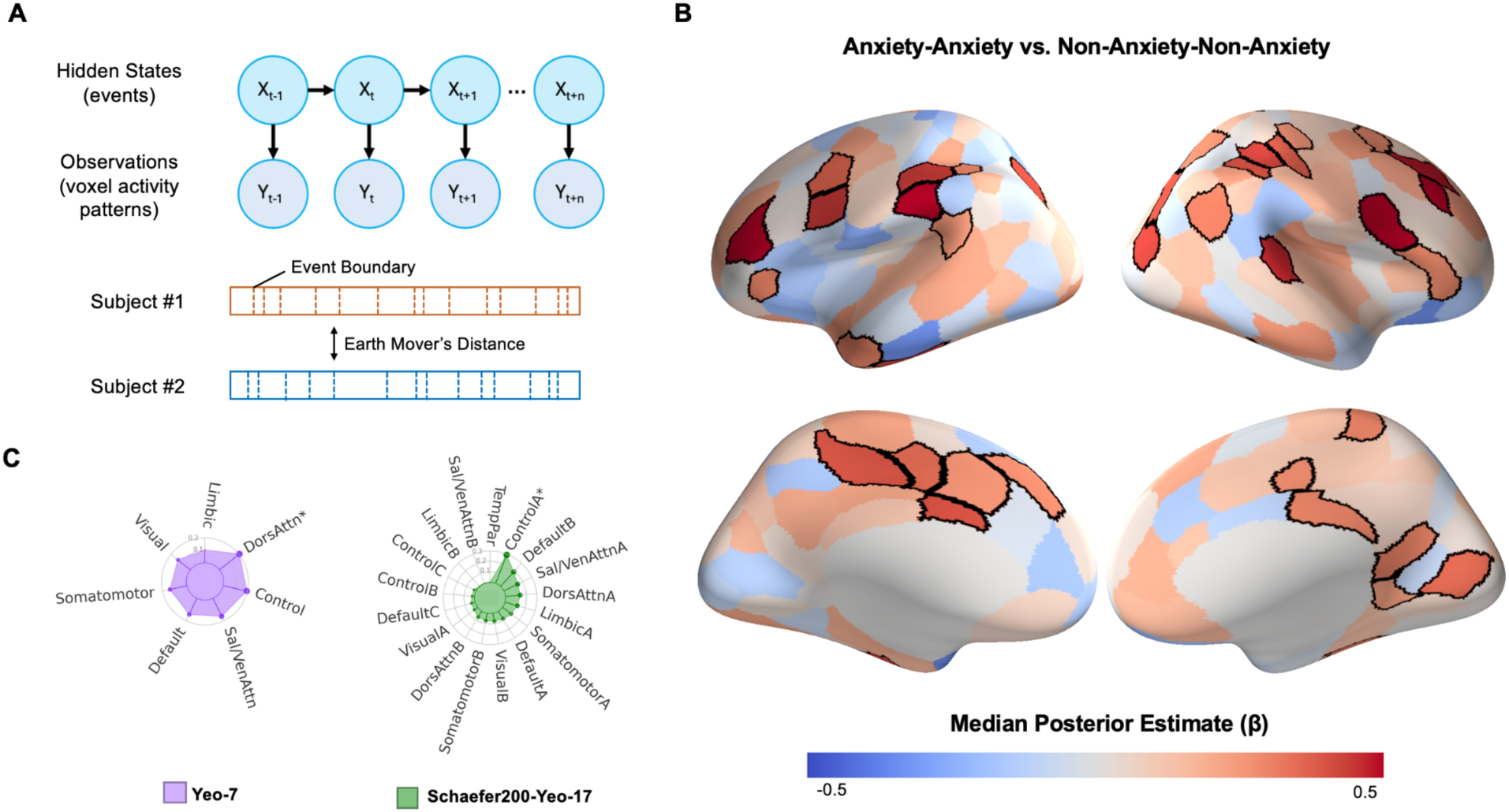
Anxiety is associated with idiosyncratic neural event boundaries. **(A)** Schematic of the Hidden Markov Model (HMM) approach for estimating individual neural event boundaries. Alignment in event boundaries between pairs of participants were quantified using Earth Mover’s Distance (EMD). **(B)** Cortical maps showing regions where anxious-anxious dyads exhibited significantly higher EMD compared to non-anxious-non-anxious dyads. Black outlines indicate regions that survive controlling for multiple comparisons at a threshold of expected PEP < 0.01. **(C)** Network overlap analysis indicated that regions with reduced alignment in anxious dyads significantly overlapped with the dorsal attention network in the Yeo-7 parcellation scheme (Yeo-7) and control network A in the Yeo-17 parcellation scheme (Schaefer200-Yeo-17). * p < 0.05.

We note that anxiety scores were more variable in the anxious group (SD = 2.35) than in the non-anxious group (SD = 1.25). Thus, an alternative explanation for our findings is that greater boundary divergence in anxious dyads may reflect larger score differences within those dyads, rather than idiosyncrasy due to trait anxiety. To address this possibility, we conducted intersubject representational analyses that relate EMD scores to a continuous measure of dyadic anxiety (see Methods). Specifically, we used Bayesian multilevel models to predict EMD scores of a dyad from their mean anxiety scores. Dyads with the highest anxiety scores would have a high mean but a small score difference. Thus, if higher mean anxiety scores tracked with higher EMD scores, it would indicate that event boundary divergence scaled with anxiety levels, rather than the difference in anxiety scores. Indeed, pairs with higher mean anxiety scores had higher EMD scores in an overlapping set of brain regions identified in Figure 2B, as well as in the ventromedial prefrontal cortex (Figure S4). No regions exhibited a credible effect in the opposite direction.

In sum, the divergence in event boundaries increased with the addition of an anxious individual to the dyad, as well as a function of the average anxiety level of the dyad, providing converging evidence that trait anxiety was associated with more individually distinct event boundaries.

### 3.3 Regions with reduced boundary alignment overlap with dorsal attention and control networks

To identify the functional networks that exhibited idiosyncratic event boundaries, we used the Network Correspondence Toolbox (Kong et al., 2025), which quantifies the overlap of a statistical map with networks defined by commonly used atlases. Overlap was computed as the Dice coefficient, with statistical significance assessed using surface-based spin permutation testing. In the Yeo 7-network parcellation scheme, regions with reduced boundary alignment in anxious-anxious dyads showed significantly greater-than-chance overlap with the dorsal attention network (Dice = 0.205, *p* = 0.014; Figure 2C). In the Yeo 17-network parcellation scheme, the regions showed significantly greater-than-chance overlap with control network A (Dice = 0.294, p = 0.001; Figure 2C), a subnetwork of the frontoparietal control network (often referred to as FPN-B in other nomenclatures; Kucyi et al., 2021) previously shown to be preferentially functionally coupled with the dorsal attention network (Dixon et al., 2018). No other canonical networks exhibited above-chance overlap.

### 3.4 Event boundaries were more idiosyncratic during anxiety provoking moments

We next asked whether this idiosyncrasy was greater during anxiety-inducing moments in the movie. We hypothesized that event boundaries would be more idiosyncratic during these moments, when appraisals of uncertainty and threat are more likely to diverge across individuals. To test this, we divided the movie into 4-TR segments and generated segment-by-segment anxiety ratings by applying a large language model (GPT-4o) to fine-grained annotations of onscreen events and dialogue (see Methods). We also used GPT-4o to generate arousal and valence ratings of the segments to include as covariates in subsequent analyses.

We first validated the GPT-4o ratings against human ratings collected in independent datasets. For anxiety, we used three films from the Emo-FilM dataset (Morgenroth et al., 2025), which contains moment-by-moment human ratings of anxiety for each film. GPT-4o anxiety ratings were correlated with human anxiety ratings in all three films (r = 0.68-0.79, all p < .001; Figure S5A; Table S2). Emo-FilM does not include arousal or valence ratings, so we validated GPT-4o arousal and valence ratings against the related dimensions of alertness and happiness in the same three films (arousal-alertness: r = 0.47-0.74; valence-happiness: r = 0.42-0.70; all p < .05; Figure S5A). For a more direct test, we additionally validated GPT-4o arousal and valence ratings against human ratings of arousal and valence for an episode of BBC’s *Sherlock* (Ke et al., 2025). Both showed strong agreement with human ratings (arousal: r = 0.77, p < .001; valence: r = 0.60, p < .001; Figure S5B). These results support the use of GPT-4o ratings as a measure of moment-by-moment affective content in our stimulus.

GPT-4o ratings were aligned with the neural event boundaries and used to predict increases in event boundary variability in the anxious group relative to the non-anxious group across ROIs (boundary variance ratio; see *Methods*), while controlling for the valence, arousal, brightness, motion, and optical flow of the corresponding segment. Average anxiety ratings of the event immediately preceding the boundary significantly predicted higher boundary variance ratio (median β = 0.252, posterior probability β > 0 = 0.978; 95% CI = [0.007, 0.495]; Figure 3B). Thus, event boundaries were more variable during moments in the movie rated as anxiety provoking, suggesting that the variability reflects meaningful, stimulus-driven differences rather than spontaneous internal fluctuations. Coefficients for the covariates are reported in Supplementary Table S3; none were credibly associated with boundary variance ratio (all posterior probabilities below 0.90).

**Figure 3.**
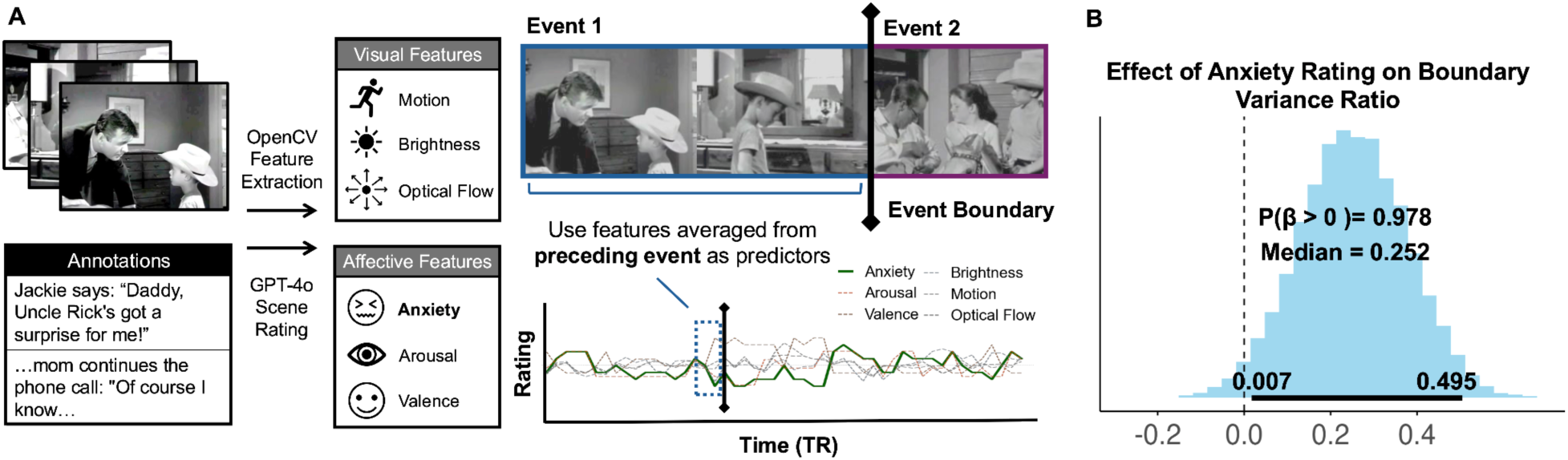
Event boundaries are more idiosyncratic during anxiety-provoking moments. **(A)** Feature extraction pipeline. The movie was annotated in 4TR segments. Scene annotations were rated by GPT-4o on anxiety, arousal, and valence while visual features (motion, brightness, optical flow) were detected from every 5 frames of the movie using OpenCV and averaged within 4TR segments. For each event boundary, features from the most recent segment preceding the boundary were used as predictors. **(B)** Posterior distribution for the effect of anxiety rating on event boundary variance ratio (variance_anxious_ / variance_control_). Higher anxiety in the preceding segment predicted greater relative variability in neural event boundaries among anxious individuals. Thick solid line indicates 95% credible interval.

### 3.5 Anxiety-provoking moments were associated with higher decoded sustained attention in both groups

Having found that boundary divergence in anxious individuals was concentrated during anxiety-inducing moments of the film, we ran a series of exploratory analyses to examine how participants attended to these moments. While the CamCAN dataset does not include behavioral measures of attention during movie-watching, we can take advantage of a previously validated neural signature of sustained attention. Specifically, the sustained attention connectome-based predictive model (saCPM; Rosenberg et al., 2016) has been shown to predict sustained attention in cognitive tasks (Rosenberg et al., 2016, 2020) and to generalize to narrative engagement during movie-watching and story-listening (Corriveau et al., 2026; Song et al., 2021). We applied the saCPM to participants’ functional data to obtain a moment-by-moment estimate of decoded sustained attention from each participant’s functional connectivity patterns (see Methods).

We defined high-anxiety and low-anxiety segments during the movie using a tertile split. Decoded sustained attention was higher during high-anxiety segments than low-anxiety segments in both groups (anxious: *t*(59) = 6.44, p < .001; non-anxious: *t*(58) = 7.27, p < .001; Figure 4). The group x segment-type interaction was not significant (*b* = -0.040, *t*(234) = -0.56, p = 0.57). Results were consistent when segment-type was defined using median and quartile splits (Figure S6), as well as when anxiety ratings of the segment were treated as a continuous variable (anxious: b = 0.14, t(59.0) = 8.15, p < .001; non-anxious: b = 0.14, t(58.0) = 7.74, p < .001; interaction: b = −0.002, t(117.0) = -0.07, p = 0.94).

**Figure 4.**
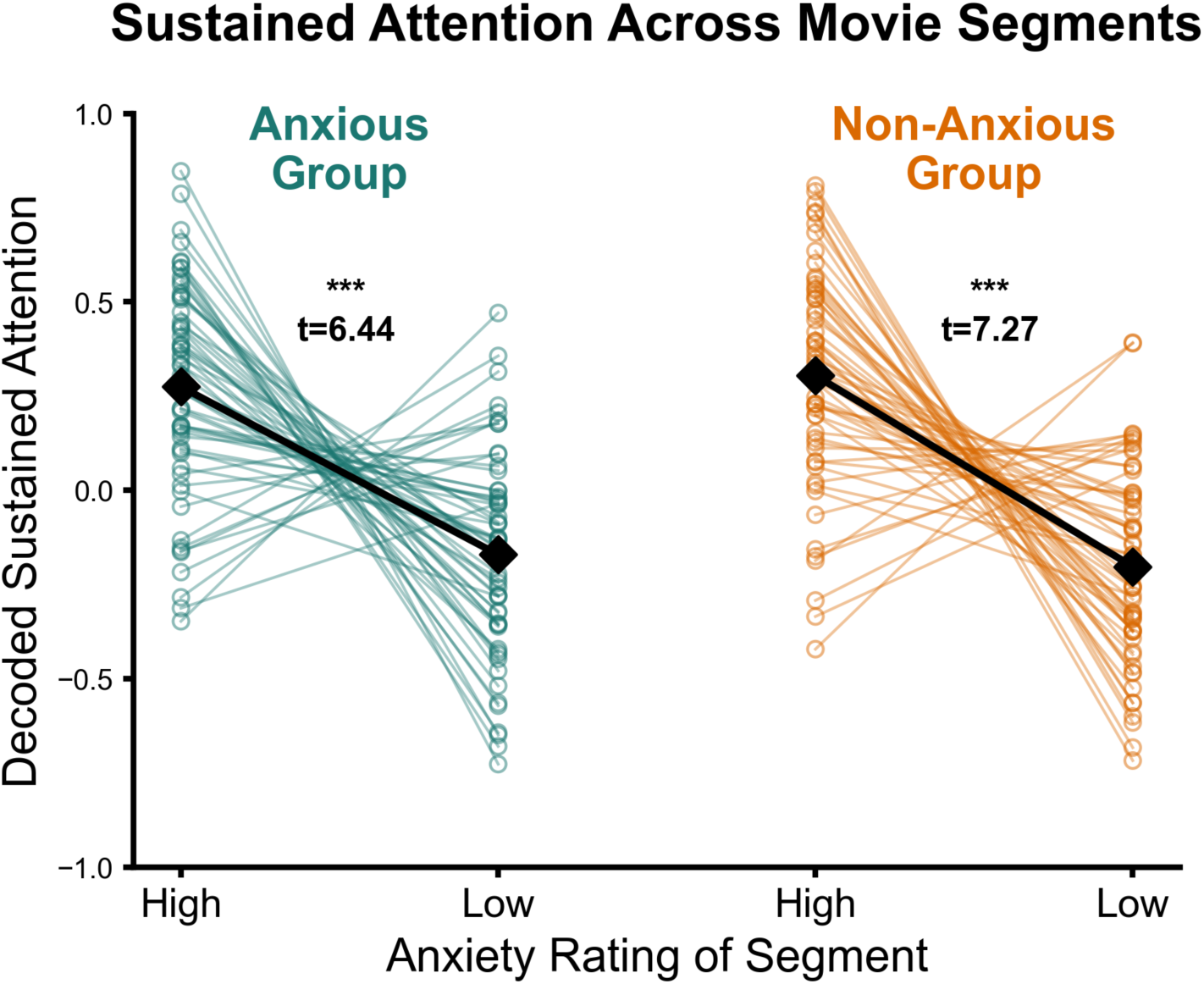
Decoded sustained attention was higher in both anxious and non-anxious individuals during high-anxiety movie moments. Each datapoint represents a single participant’s decoded sustained attention score during high-versus low-anxiety segments of the film. Segments were grouped based on a tertile split. Solid black diamonds indicate group means. No group interaction on the difference between high-anxiety vs. low-anxiety moments was observed.

Together, these results suggest that anxiety-provoking moments of the film elicited heightened sustained attention. This effect was present in both anxious and non-anxious participants, suggesting that the divergent event boundaries in the anxious group are not attributable to anxious participants disengaging from the stimulus during high-anxiety moments.

## 4. Discussion

We applied data-driven state-detection methods to movie-watching fMRI data to investigate the relationship between trait anxiety and neural event segmentation. Higher trait anxiety was associated with idiosyncratic neural event boundaries, particularly in the dorsal attention and frontoparietal control networks. The divergence in boundary alignment between the two groups was greatest during moments rated as more anxiety-provoking, indicating that the variability in boundary alignment fluctuated with narrative content. Together, our results suggest that trait anxiety alters how the brain represents and organizes continuous experience.

Our findings parallel a recent study finding that anxious individuals showed unique alterations along different locations of the amygdala-ventral PFC white matter tract, whereas low trait-anxiety individuals displayed convergent morphology (Kim & Kim, 2022). Thus, anxiety may be characterized not by a single, shared neural signature but rather by diverse and individualized patterns of brain activity and morphology. This pattern aligns with a broader phenomenon where individuals with adaptive traits exhibit similar neural responses, while individuals with maladaptive traits differ not only from healthy controls but also from others with the same traits (Berman et al., 2013; Byrge et al., 2015; J. Chen et al., 2026). For example, lonely individuals have individually idiosyncratic neural representations of movies and cultural concepts, while non-lonely individuals have shared representations (Baek et al., 2023; Broom et al., 2024). In a similar vein, pessimists exhibit divergent neural responses during future thinking but optimists converge on similar patterns (Yanagisawa et al., 2025), and individuals with worse reading ability have more idiosyncratic responses than better readers (Jangraw et al., 2023).

This phenomenon has been termed the Anna Karenina effect, after Tolstoy’s observation that “all happy families are alike; each unhappy family is unhappy in its own way”. In the context of anxiety, the idiosyncrasy in neural event segmentation may stem from the heterogeneity in the underlying etiology: anxious individuals may have different biases in attending to, interpreting, and predicting unfolding events while building and updating event models. For example, some individuals may be primarily concerned with monitoring and preventing errors in routine, whereas others may devote attention to monitoring novel threats in uncertain situations, resulting in individualized patterns of attention and interpretation over time (Roemer et al., 1997; Watkins, 2008).

Interestingly, a recent study found seemingly discrepant results, where high anxiety individuals behaviorally segmented movies more consistently than non-anxious individuals when asked to explicitly divide a movie into events (Bein et al., 2025). The authors interpreted this consistency as reflecting anxious individuals’ preference for predictable and stable environments. We speculate that the contrasting results may arise from Bein and colleagues intentionally selecting a movie that did not evoke strong emotions, while our study used a suspenseful film. Thus, one possibility is that anxious individuals’ preference for stability promotes consistency during emotionally neutral narratives, but this preference is challenged during suspenseful narratives, increasing person-specific appraisals of uncertainty and producing idiosyncratic event boundaries. Another possibility is that anxiety may increase consistency during explicit behavioral segmentation by encouraging reliance on predictable cues such as scene cuts, even if neural event transitions diverge due to idiosyncratic representational shifts. Future work combining behavioral and neural measures within the same stimulus will be needed to distinguish these accounts.

Our findings contribute to growing evidence that stable individual differences modulate neural responses to ambiguous, naturalistic stimuli. Previous work has demonstrated that traits including paranoia, hostile attribution bias, and personality dimensions influence inter-subject neural synchrony during narrative perception (Finn et al., 2018; Matz et al., 2022; Lyu et al., 2024). Paralleling our results, a recent study found that loneliness was associated with idiosyncratic neural event boundaries in the default mode network (DMN) (Lu et al., 2025). Together, this emerging literature suggests that stable individual traits not only shape the degree of neural synchrony during shared experiences, but also the temporal structure by which individuals parse ongoing events.

While anxious individuals also exhibited idiosyncratic neural event boundaries in DMN regions, including the dorsomedial prefrontal cortex and precuneus, the set of identified regions overlapped most prominently with the dorsal attention network (DAN), a network implicated in the top-down allocation of visuospatial attention (Corbetta & Shulman, 2002; Ungerleider, 2000). When the networks were examined at a more granular level, the regions showed significant overlap with Control Network A, a subnetwork of the frontoparietal control network (FPN) that preferentially couples with the DAN. In contrast to Control Network B, which is more strongly coupled with the DMN and is thought to support control of internally directed cognition, Control Network A has been associated with the control of perceptual attention during externally directed cognition (Dixon et al., 2018; Waskom et al., 2014). Similar to anxiety-related attentional biases observed in other domains (Eysenck et al., 2007; Mathews & MacLeod, 2005), anxious individuals may adopt person-specific attentional priorities when parsing anxiety-provoking moments in a narrative. Consistent with the idea that anxious individuals vary in how they infer latent states under threat, prior behavioral work found that anxious individuals differ in how they update latent task states when rules change during aversive learning (Zika et al., 2023).

A potential alternative explanation for higher boundary variability during anxiety-inducing moments in anxious individuals is that these participants disengaged from the stimulus during such moments, producing more variable neural activity through off-task processing rather than through idiosyncratic stimulus-driven segmentation. Our analyses of decoded sustained attention provide evidence against this account. Using a neural index of sustained attention, we found that anxiety-provoking moments of the film elicited heightened sustained attention in both anxious and non-anxious participants, suggesting that anxious individuals did not disengage during these moments. This finding is consistent with prior work showing that emotionally salient and threat-related content preferentially captures and sustains attention (Koster et al., 2004; Pourtois et al., 2013). We note that the saCPM provides a global index of sustained attention (Corriveau et al., 2025), and would not detect differences in which aspects of the content anxious individuals attended to. Indeed, we believe such differences could contribute to the observed boundary divergence.

Although our study did not include behavioral measures of memory, the patterns we observed raise the question of how idiosyncratic event boundaries might shape downstream memory for shared experiences. Event boundaries are well-established to influence how continuous experience is encoded, structured, and retrieved (Clewett et al., 2019; DuBrow & Davachi, 2013; Michelmann et al., 2023; Silva et al., 2019; Swallow et al., 2009; Yates et al., 2023). Prior work has shown that individuals whose neural event boundaries are more aligned during a narrative subsequently exhibit more similar recall and interpretations of that narrative (Sava-Segal et al., 2023). Extending this to our findings, the idiosyncratic boundaries we observed in anxious individuals suggest that anxious individuals may form memories of shared experiences that are idiosyncratic, both relative to non-anxious individuals and to other anxious individuals. Future work combining neural event segmentation with assessments of memory, comprehension, and affective responses could directly test this prediction and clarify whether idiosyncratic boundaries contribute to the memory differences observed in anxiety.

Our study utilized a large-scale publicly available fMRI dataset and data-driven methods to infer neural event boundaries without disrupting ongoing cognition, examining event segmentation as it occurs during naturalistic viewing conditions. Additionally, we implemented two methodological innovations. First, we quantified boundary similarity using Earth Mover’s Distance (EMD), which provides a more fine-grained measure of temporal alignment than the previously adopted boundary-matching approaches by considering the temporal proximity of event boundaries between individuals. Second, we employed large language models and computer vision tools to systematically characterize stimulus features across movie scenes, allowing us to directly test whether boundary variability in anxious individuals tracked content characteristics rather than reflecting differences in nonspecific fluctuations of neural patterns.

Several limitations of our study should be acknowledged. First, our measure of anxiety, HADS-A, does not differentiate between anxiety subtypes such as generalized, social, or panic-related anxiety, leaving open the question of whether the patterns observed here are common across anxiety dimensions or specific to particular subtypes. Second, our groups were defined by HADS-A scores rather than clinical diagnosis, which may limit generalizability to clinical populations. Third, our study did not include behavioral measures of memory or comprehension of the stimulus, leaving the behavioral consequences of idiosyncratic event boundaries to be established in future work. Future investigations using our approach with comprehensive anxiety assessments, behavioral measures, varied stimuli, and clinical populations will clarify how neural event segmentation varies across anxiety dimensions and relates to clinical phenomena.

In conclusion, these results shed new light on how trait anxiety modulates neural processing of dynamic, continuous experiences. The finding that anxiety is associated with more idiosyncratic event boundaries within the DAN and FPN underscores the critical role of top-down attentional mechanisms in shaping individual event representations. Our approach establishes a neurocomputational framework for understanding anxiety-related alterations in the temporal organization of dynamic information. More broadly, the analytical framework developed here provides a generalizable approach for investigating the influence of other traits and clinical conditions on event cognition, revealing the individual differences in how people impose structure on the stream of experience.

## Data and Code Availability

Code used in analysis is available at https://github.com/20-alicial/trait-anxiety-event-seg. The Cam-CAN dataset from (Shafto et al., 2014; Taylor et al., 2017) is available upon request at the CBU Data Portal: https://opendata.mrc-cbu.cam.ac.uk/projects/camcan/

## Author Contributions

ATHL: Conceptualization; Data curation; Methodology; Formal analysis; Project administration; Software; Visualization; Writing - original draft; and Writing - review & editing. JC: Methodology; Software; and Writing - review & editing. JSP: Methodology; Software. HK: Data curation. YCL: Conceptualization; Data curation; Methodology; Project administration; Supervision; Writing - original draft; and Writing - review & editing

## Acknowledgements

Cam-CAN data collection was funded by the UK Biotechnology and Biological Sciences Research Council (grant number BB/H008217/1) and the UK Medical Research Council and University of Cambridge.

## Supplementary Information

### Supplementary Materials

**Figure S1:**
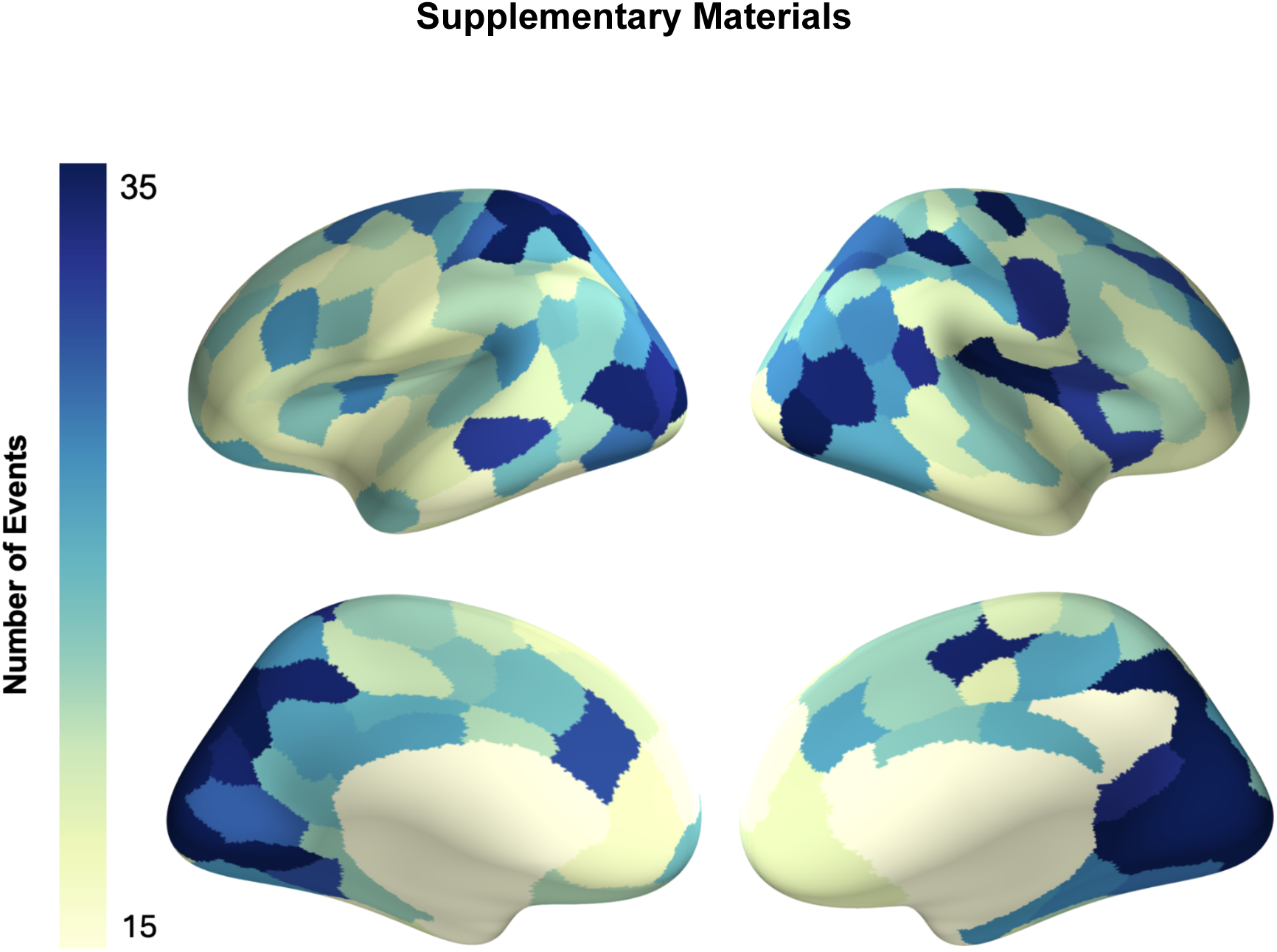
Number of events identified in each ROI when GSBS is fit to all participants. The analysis revealed a similar posterior-to-anterior gradient in event segmentation rates that was observed when GSBS was fit separately for anxious and non-anxious participants.

**Figure S2.**
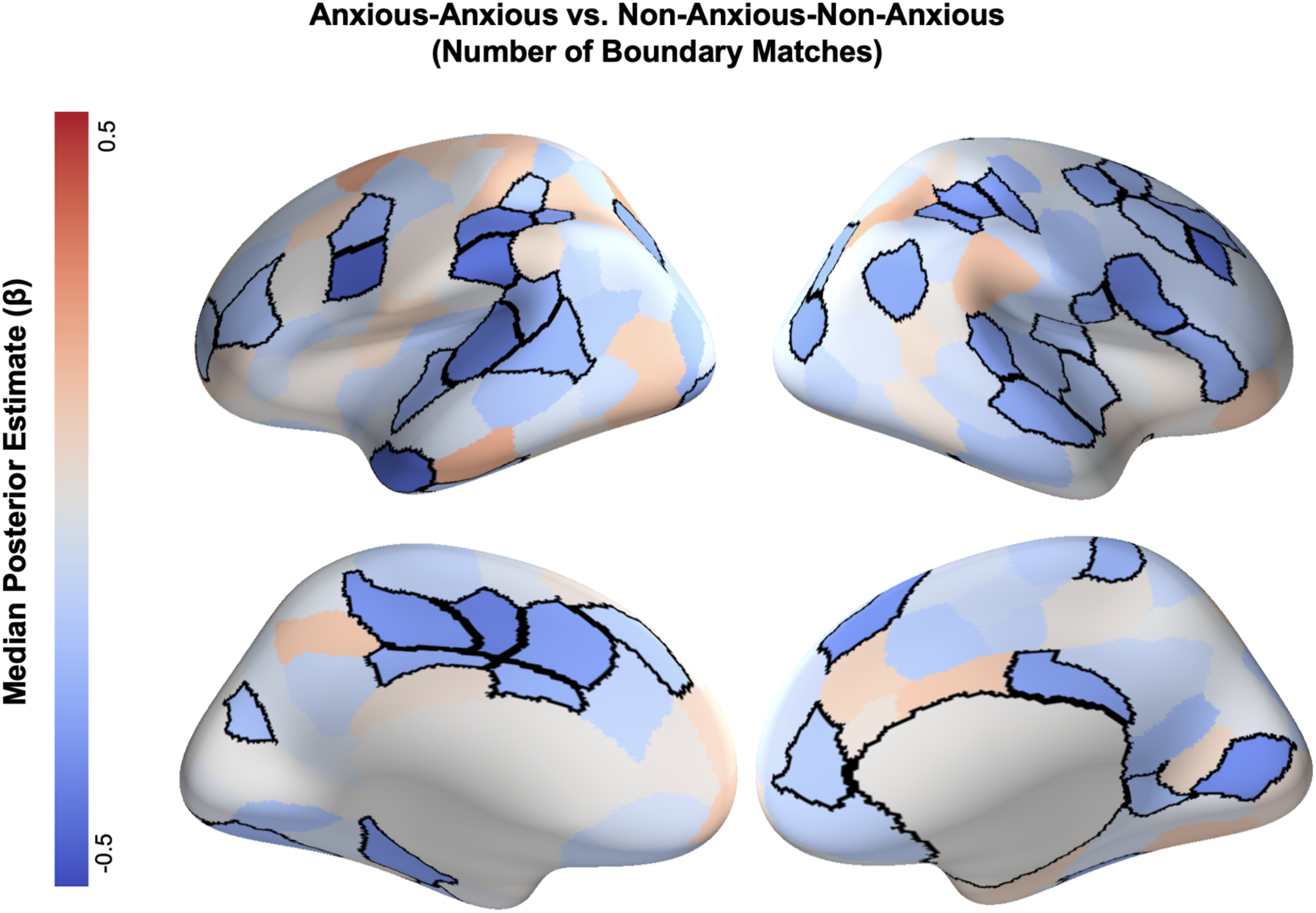
Brain regions where boundary alignment is lower for anxious dyads than non-anxious dyads. To assess whether our findings depended on the choice of boundary alignment metric, we repeated the analyses with alignment assessed using the number of boundary matches within 1TR. The analyses identified a similar set of regions where event boundaries were less aligned between anxious individuals than between non-anxious individuals. Note that the effect is negative here as lower boundary alignment is indicated by fewer boundary matches, as opposed to higher EMD.

**Figure S3.**
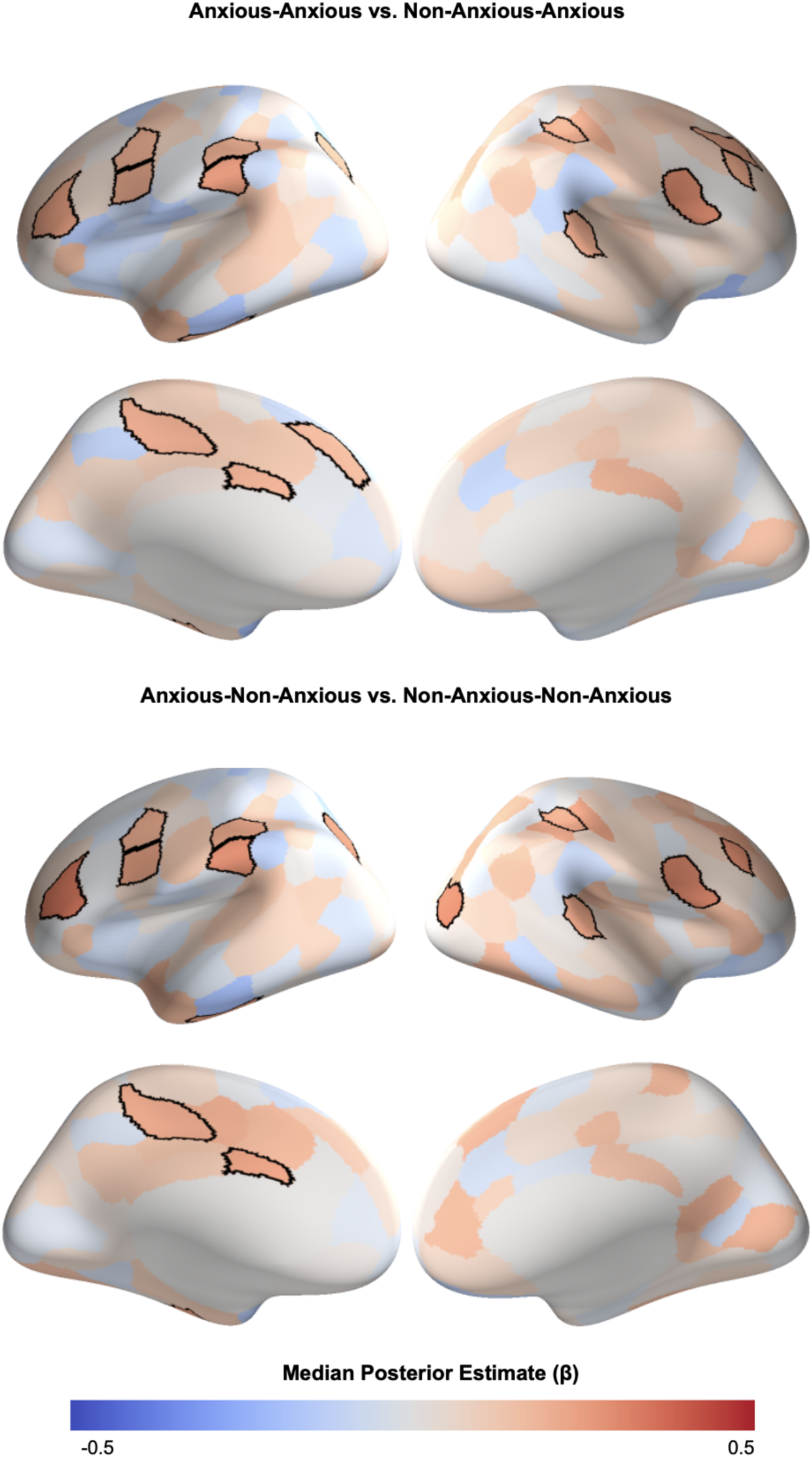
Brain regions where boundary alignment is lower for anxious dyads than mixed dyads (top) and lower for mixed dyads than non-anxious dyads (bottom). These results indicate that alignment in event boundaries decreased with an additional anxious individual in the dyad.

**Figure S4.**
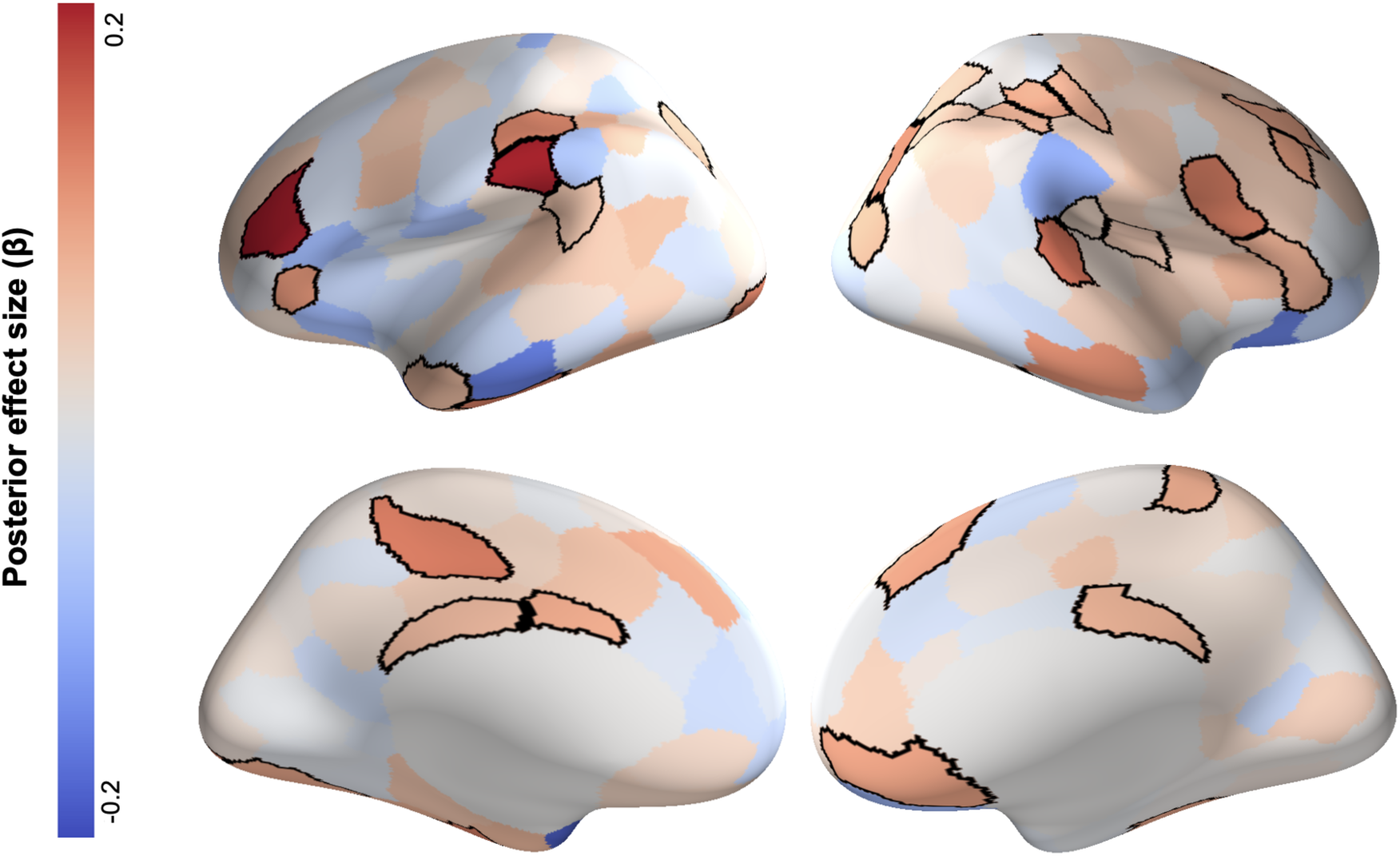
Intersubject representational analysis between mean dyadic anxiety score and EMD. We used Bayesian multilevel models to predict EMD from the average anxiety score of a dyad. Dyads with higher average anxiety scores had higher EMD scores, indicating that their event boundaries were less aligned.

**Figure S5.**
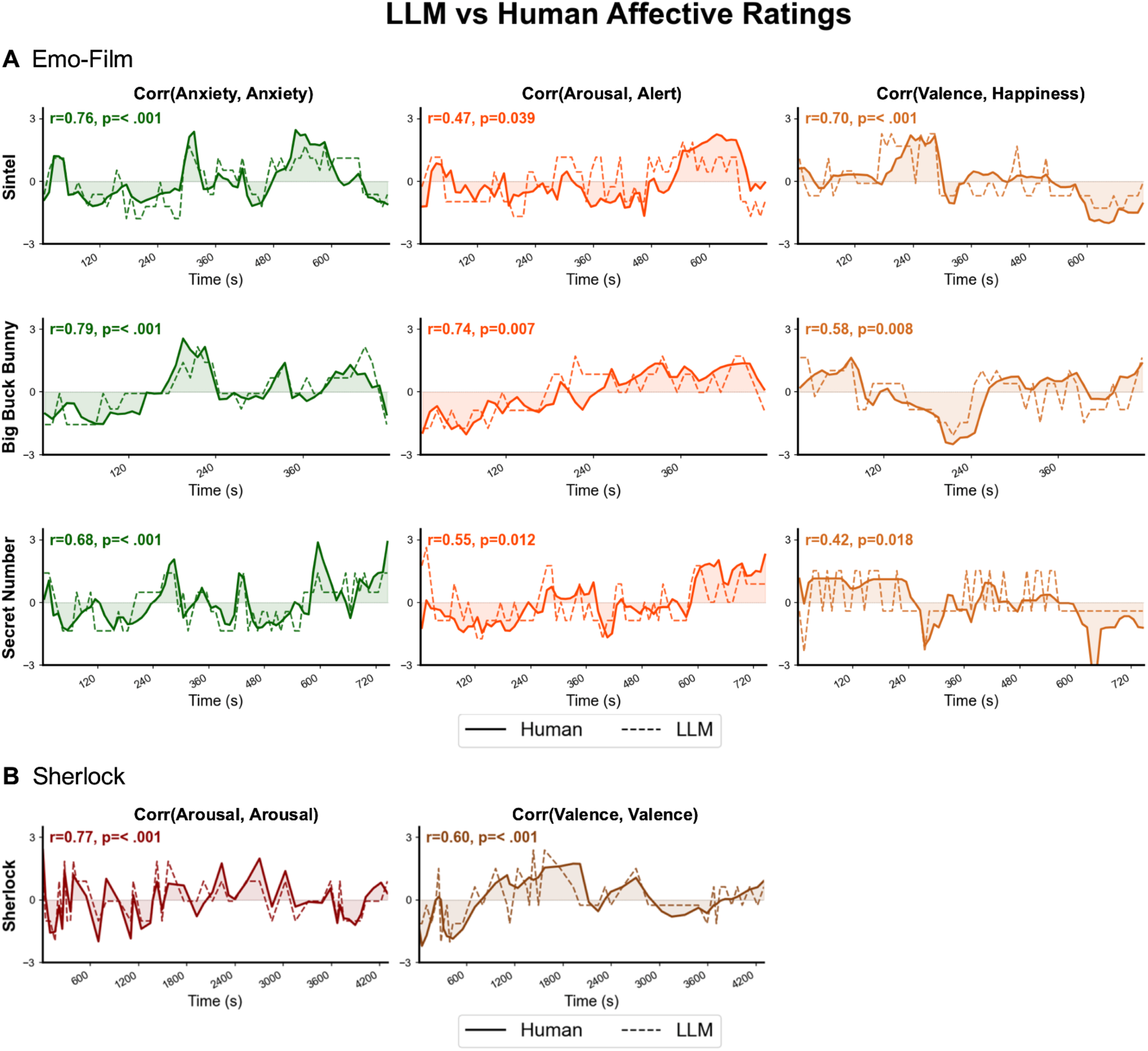
GPT-4o affective ratings were comparable to human consensus ratings across four films. Solid lines indicate human ratings; dashed lines indicate GPT-4o ratings. (A) For Emo-FilM stimuli *Sintel*, *Big Buck Bunny*, and *The Secret Number*, GPT-4o anxiety ratings are compared to participant anxiety ratings, GPT-4o arousal ratings to participant alertness ratings, and GPT-4o valence ratings to participant happiness ratings. For Sherlock, GPT-4o arousal and valence ratings are compared to participant arousal and valence ratings. Pearson correlations and p-values are derived from phase-randomization shuffling the 5,000 surrogate time series.

**Figure S6.**
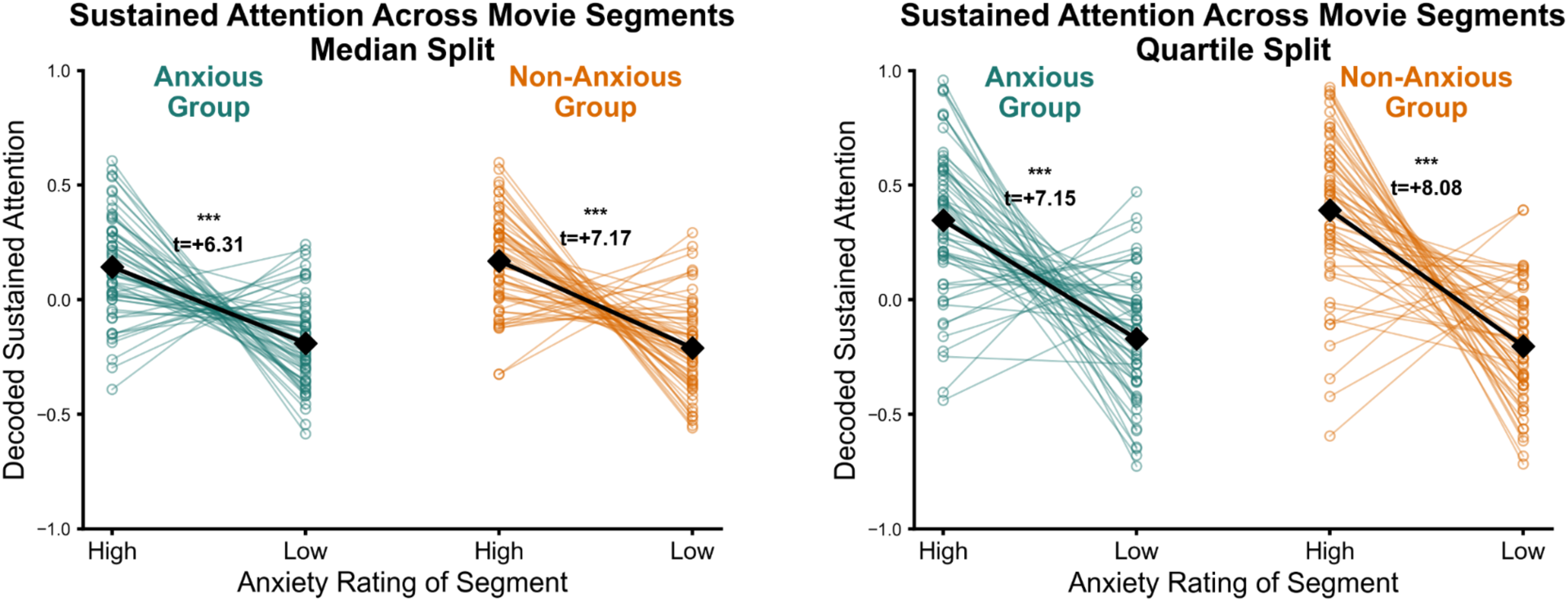
Sustained attention results replicated across median and quartile anxiety-rating splits. Both anxious and non-anxious participants showed significantly greater sustained attention during high-anxiety compared to low-anxiety moments. **(Left)** Median split.. **(Right)** Quartile split.

**Table S1.**
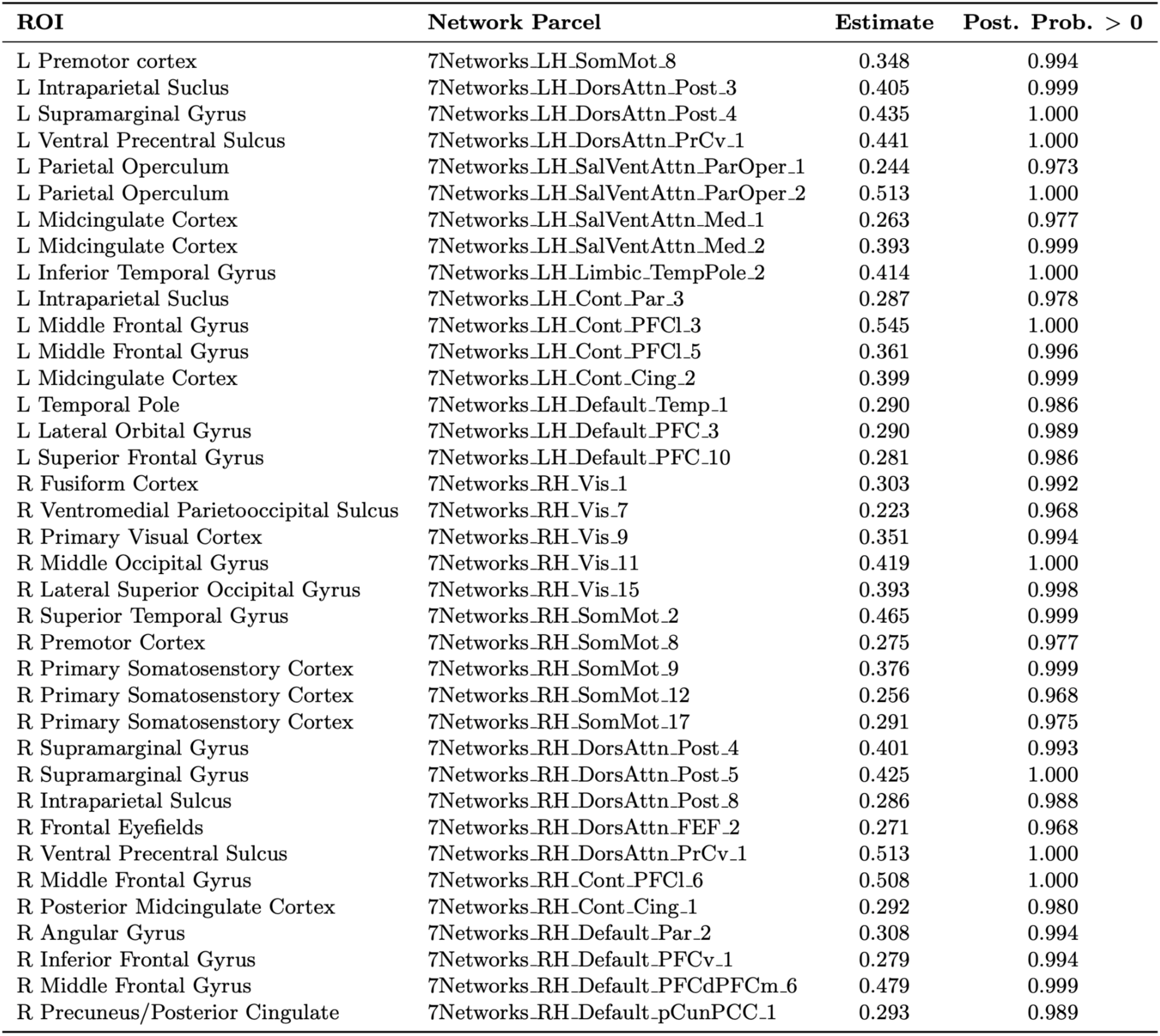
Full list of 37 ROIs exhibiting higher EMD in anxious-anxious versus non-anxious-non-anxious dyads that survive controlling for multiple comparisons at PEP < 0.01. The table displays the posterior mean estimate and posterior probabilities of a positive effect of anxiety on dyadic EMD.

**Table S2.** Validation results of GPT-4o affective rating pipeline. Pearson correlation coefficients between GPT-4o ratings and human consensus ratings across four films and five dimension pairs from two independent datasets. P-values were derived from phase-randomization permutation tests with 5,000 surrogate time series.

| Film | GPT → Human Dimension Pair | *r* | *p* |
| --- | --- | --- | --- |
| Sintel | Anxiety → Anxiety | 0.763 | < 0.001 |
|  | Arousal → Alert | 0.469 | 0.039 |
|  | Valence → Happiness | 0.698 | 0.001 |
| Big Buck Bunny | Anxiety → Anxiety | 0.794 | 0.001 |
|  | Arousal → Alert | 0.743 | 0.007 |
|  | Valence → Happiness | 0.581 | 0.008 |
| The Secret Number | Anxiety → Anxiety | 0.677 | < 0.001 |
|  | Arousal → Alert | 0.545 | 0.012 |
|  | Valence → Happiness | 0.420 | 0.018 |
| Sherlock | Arousal → Arousal | 0.774 | < 0.001 |
|  | Valence → Valence | 0.604 | < 0.001 |

**Table S3.** Covariate coefficients from the Bayesian multilevel model predicting boundary variance ratio with moment-by-moment movie features. Median posterior estimates, 95% credible intervals, and posterior probabilities (probability that β > 0 or β < 0, whichever is larger) are reported for each covariate.

| Covariate | Median $\beta$ | 95% CI | Posterior Probability |
| --- | --- | --- | --- |
| Arousal | −0.083 | [−0.293, 0.126] | 0.779 |
| Valence | 0.118 | [−0.068, 0.306] | 0.891 |
| Brightness | −0.003 | [−0.083, 0.077] | 0.525 |
| Motion | −0.124 | [−0.335, 0.079] | 0.888 |
| Optical Flow | 0.067 | [−0.120, 0.264] | 0.754 |

## Large Language Model Prompt

### System Prompt

You are an expert movie rater, rating scenes from a movie as it unfolds.

### User Prompt

Below is everything the viewer has already seen, in order. Do NOT use any additional information:

<<< BEGIN SEEN SCENES >>>

{All previous scene annotations concatenated chronologically}

<<< END SEEN SCENES >>>

Now here is the CURRENT scene:

{Current scene annotation}

Rate the {dimension_name} level of ONLY the CURRENT scene on a scale of 1-7, using the definition below.

Return a valid JSON like this:

{{"{dimension_name}":2}}

{dimension_name} definition:

{dimension_name} – {dimension_def}

### Affective Dimension Definitions

- Arousal refers to feeling very mentally or physiologically alert, activated and/or energized, with 1 being drowsy and sluggish and 7 being highly alert and energized
- Valence refers to how positive or negative the emotional experience feels, with 1 being very unpleasant and negative and 7 being very pleasant and positive
- Anxiety refers to feeling tense, nervous, or apprehensive about what may happen, with 1 being calm and at ease and 7 being worried and on edge

Large language model prompt template. Each scene was rated independently on each affective dimension (arousal, valence, anxiety) using GPT-4o (temperature = 0). The model received a system prompt establishing its role as an expert movie rater, and a user prompt containing all previously seen scene annotations as rolling context, the current scene annotation, and the definition for that single dimension. The three dimensions were rated in separate passes over the full sequence of scenes, so that ratings on one dimension could not influence ratings on another.

